# Bioconversion of Glucose-Rich Lignocellulosic Wood Hydrolysates to 3-Hydroxypropionic Acid and Succinic Acid using Engineered *Saccharomyces cerevisiae*

**DOI:** 10.1101/2023.11.23.568477

**Authors:** Scott Bottoms, Christina Mürk, Huadong Peng, Rodrigo Ledesma-Amaro, Mart Loog

## Abstract

**Background:** 3-hydroxypropionic acid (3-HP) and succinic acid (SA) were announced as two of the top twelve value-added platform chemicals from biomass out of a group of over 300 potential compounds that could be made from biomass in a government report in 2004 (Werpy and Petersen, 2004) and in an updated report in 2010 (Bozell and Petersen, 2010). The screening criteria used in the report classified 3-HP and SA as direct petroleum replacement building block chemicals. 3-HP is a precursor to several high-value compounds, such as acrylic acid, 1,3-propanediol, acrylamide, and methyl acrylates, that ultimately end up in products such as fibers, contact lenses, diapers, fabric coatings, and other super absorbent polymers (SAPs). SA is a high-value platform chemical used in polyester production and a precursor for nylon and other bioplastics. Additionally, these reports identified pathways to building block compounds from sugars. Yeast fermentations were identified in these reports as a preferred potential pathway to 3-HP and SA production from sugars because of yeasts’ natural low pH tolerance.

**Results:** The laboratory strain *Saccharomyces cerevisiae* BY4741 was engineered to produce either 3-HP or SA. These yeasts can convert fermentable sugars from glucose-rich lignocellulosic hardwood feedstocks into organic acid products such as 3-HP and SA under low pH conditions using exponential fed-batch cultivation strategies. Glucose-rich wood sugars provided a better growth environment for the engineered yeast strains, increasing production titers by 6.1 and 6.5 times for SA and 3-HP, respectively.

**Conclusions:** This study shows the potential of locally produced glucose-rich wood sugars to increase the production of platform chemicals necessary in the production of biobased polymers by engineered yeast cell factories.

## Introduction

3-hydroxypropionic acid (3-HP) and succinic acid (SA) were announced as two of the top twelve value-added platform chemicals from biomass out of a group of over 300 potential compounds that could be made from biomass in a government report in 2004 (Werpy and Petersen, 2004) and in an updated report in 2010 (Bozell and Petersen, 2010). The screening criteria used in the report classified 3-HP and SA as direct petroleum replacement building block chemicals. 3-HP is a precursor to several high-value compounds, such as acrylic acid, 1,3-propanediol, acrylamide, and methyl acrylates, that ultimately end up in products such as fibers, contact lenses, diapers, fabric coatings, and other super absorbent polymers (SAPs). SA is a high-value platform chemical used in polyesters and a precursor for nylon and other bioplastics. Additionally, these reports identified pathways to building block compounds from sugars. Yeast fermentations were identified in these reports as a preferred potential pathway to valuable organic acids from sugars because of yeasts’ natural low pH tolerance. Conveniently, 3-HP has a pKa of 4.87 and SA a pKa_1/2_ of 4.2 and 5.6; thus, utilizing a fermentation process at low pH improves the overall production economics compared to neutral or high pH production microbes.

*Saccharomyces cerevisiae* BY4741-3HP was engineered to produce 3-HP through the β-alanine biosynthetic pathway (Sun et al., 2023) as it was identified as the most economically attractive according to a biosynthetic metabolic model (Borodina et al., 2015) as β-alanine (3-aminopropionic acid) is the only occurring β-type amino acid and is an abundant microbial metabolite as it is a precursor of pantothenic acid which is an intermediate metabolite in coenzyme A and acyl carrier protein (Wang et al., 2021).

## Methods

### Strains and Chemicals

*S. cerevisiae* BY4741 (*MATa his3Δ1 leu2Δ0 met15Δ0 ura3Δ0*) was utilized in all experiments, as described by Shaw et al. (2022). The yeast strains were made prototrophic using single-copy plasmids for auxotrophic coverage, as outlined by Mülleder et al. (2016). The 3-HP standard was obtained from Tokyo Chemical Industry Company (TCI), the SA standard from Acros Organics (Belgium), D-Glucose anhydrate from Research Products International (DPI, Prospect, IL, USA), and C6-rich wood sugars from Fibenol OÜ (Imavere, Estonia).

### Yeast Growth

Yeast cells were cultured from glycerol stocks in standard complex medium (YPD) at 30 °C, with solid medium containing 10 g/L tryptone, 5 g/L yeast extract, 10 g/L agar, and 20 g/L D-glucose. Liquid cultures without agar were shaken at 250 RPM under the same conditions.

### Growth Medium

Synthetic Mineral Medium (SMM) comprised 7.5 g/L (NH_4_)_2_SO_4_, 14.4 g/L KH_2_PO_4_, 0.9 g/L MgSO_4_ · 7H_2_0, with trace elements and vitamin solutions. The trace elements solution contains (per L) 9 g CaCl_2_.2H_2_O, 9 g ZnSO_4_.7H_2_O, 6 g FeSO_4_.7H_2_O, 2 g H_3_BO_3_, 2 g MnCl_2_.4H_2_O, 0.8 g Na_2_MoO_4_.2H_2_O, 0.6 g CoCl_2_.6H_2_O, 0.2 g CuSO_4_.5H_2_O, 0.2 g KI, and 30 g EDTA. The vitamin solution contains (per L) 50 mg biotin, 200 mg *p*-aminobenzoic acid, 1 g nicotinic acid, 1 g calcium pantothenate, 1 g pyridoxine HCl, 1 g thiamine HCl, and 25 g myo-inositol. SMD and SMW denoted medium combined with either glucose or wood sugars, respectively.

### Cultivation Conditions

Growth morphologies were visualized in 96-well microtiter plates and 250 mL flasks with working volumes of 200 μL and 50 mL of SMD(2%) and SMW(2%), agitated at 1200 RPM or 250 RPM, respectively. Optical density values of microtiter plates were measured at 600 nm (BioTek Synergy MX), and culture flasks were measured using a spectrophotometer (Thermo Genesys 50) in standard polystyrene 1 mL cuvettes.

### Seed Preparation for Bioreactors

Freshly grown colonies from a YPD plate were used to inoculate SMD(2%), shaken at 250 RPM at 30 °C overnight. This culture was split into two flasks with fresh SMD(2%) the following morning and continued shaking for approximately two doubling times. The two flasks were combined and centrifuged, and the pellet was resuspended in fresh SMD(2%) for bioreactor seeding.

### Fed-batch Cultivations

Cultivations were carried out in 1-L bioreactors (Applikon Biotechnology, Delft, Netherlands) with an initial batch volume of 300 mL at pH 5.0 and 30 °C. Dissolved oxygen was maintained at 25% or higher using 1-vvm aeration and agitation speeds between 500 to 950 RPM. All experiments were seeded at OD600 of 0.5. The composition of the cultivation off-gas (CO_2_/O_2_) was measured using an inline gas analyzer (BlueSens gas sensor GmbH, Herten, Germany). Data was recorded and curated with BioXpert v2.95 (Applikon Biotechnology, Delft, Netherlands). Batch growth medium included 20 g/L D-glucose (SMD(2%)), while fed-batch medium had higher growth salts concentrations, comprising 148 g/L (NH_4_)_2_SO_4_, 19.7 g/L KH_2_PO_4_, and 7.8 g/L MgSO_4_ · 7H_2_0, 8 mL trace elements solution, and 5 mL vitamin solution at pH 5.0 with 0.005% antifoam 204 (Sigma) added in the feed medium only. Either 300 g/L of D-glucose or 300 g/L of glucose-rich wood sugars were added to the concentrated growth salts.

The wood sugars were centrifuged at 15,000 x g for 10 minutes before being added to the concentrated growth salts to remove unsuspended solids. Unsuspended solids would clog the feed tubing in the current configuration. In the case of SA fed-batch production, 1000 ng/μL of tetracycline A was added to the batch and feed medium as previously (Shaw et al., 2022). Initial carbon-limited feeding rates were started at 0.15 1/h. Periodic culture samples were taken and measured for optical densities and LC analysis. LC samples were filtered through a 0.45-micron syringe filter and stored at -20 °C until analysis.

### HPLC Analysis

For monosaccharide detection, 20 μL of the sample was injected into a Shimadzu LC-2050C 3D Plus system equipped with a Shimadzu RID-20A refractive index detector and an Aminex HPX-87P LC column (300 X 7.8 mm) at 85 °C using a mobile phase of MilliQ H2O at a flow rate of 0.6 mL/min for 40 minutes. Organic acid detection analyses were performed using the same system equipped with an Aminex HPX-87H LC column (300 X 7.8 mm) at 60 °C with an H^+^ precolumn. The mobile phase for organic acid detection was 1 mM H2SO4 at a flow rate of 0.6 mL/min for 45 minutes. Note that an H^+^/CO^3-^ deashing precolumn is attached inline during monosaccharide detection runs but not stored in the column oven as the maximum operating temperature is 60 °C.

## Results

Engineered *S. cerevisiae* BY4741 can convert fermentable sugars from glucose-rich lignocellulosic hardwood biomass feedstocks into platform chemical products such as 3-HP and SA under low pH conditions (Table 1). The tested yeast strains were able to produce 14.63 ± 0.29 g/L of SA by feeding wood sugars (*n=3*) and 2.39 ± 0.093 g/L of SA by feeding D-glucose (*n=3*) which is 6.1 times improvement in SA titers (Fig. 2). Concurrently, engineered yeasts produced 9.71 ± 0.168 g/L of 3-HP by feeding wood sugars (*n=2*) and 1.52 ± 0.006 g/L of 3-HP with D-glucose (*n=3*), a 6.4 times improvement in 3-HP titers (Fig. 1).

**Table 1.** Organic acid titers for *S. cerevisiae* BY4741 using different carbon sources in fed-batch.

| Strain | D-glucose Feed |  | Wood Sugar Feed |  |
| --- | --- | --- | --- | --- |
|  | 3-HP (g/L) | SA (g/L) | 3-HP (g/L) | SA (g/L) |
| BY4741 parental | 0.00 ± 0.00 | 0.09 ± 0.03 | 0.14 ± 0.04 | 0.07 ± 0.07 |
| BY4741-3HP (SHP413) | 1.52 ± 0.01 | 0.00 ± 0.00 | 9.71 ± 0.17 | 0.24 ± 0.12 |
| BR4741-SA (YLS108-2) | 0.09 ± 0.08 | 2.39 ± 0.09 | 0.00 ± 0.00 | 14.68 ± 0.29 |

**Figure 1.**
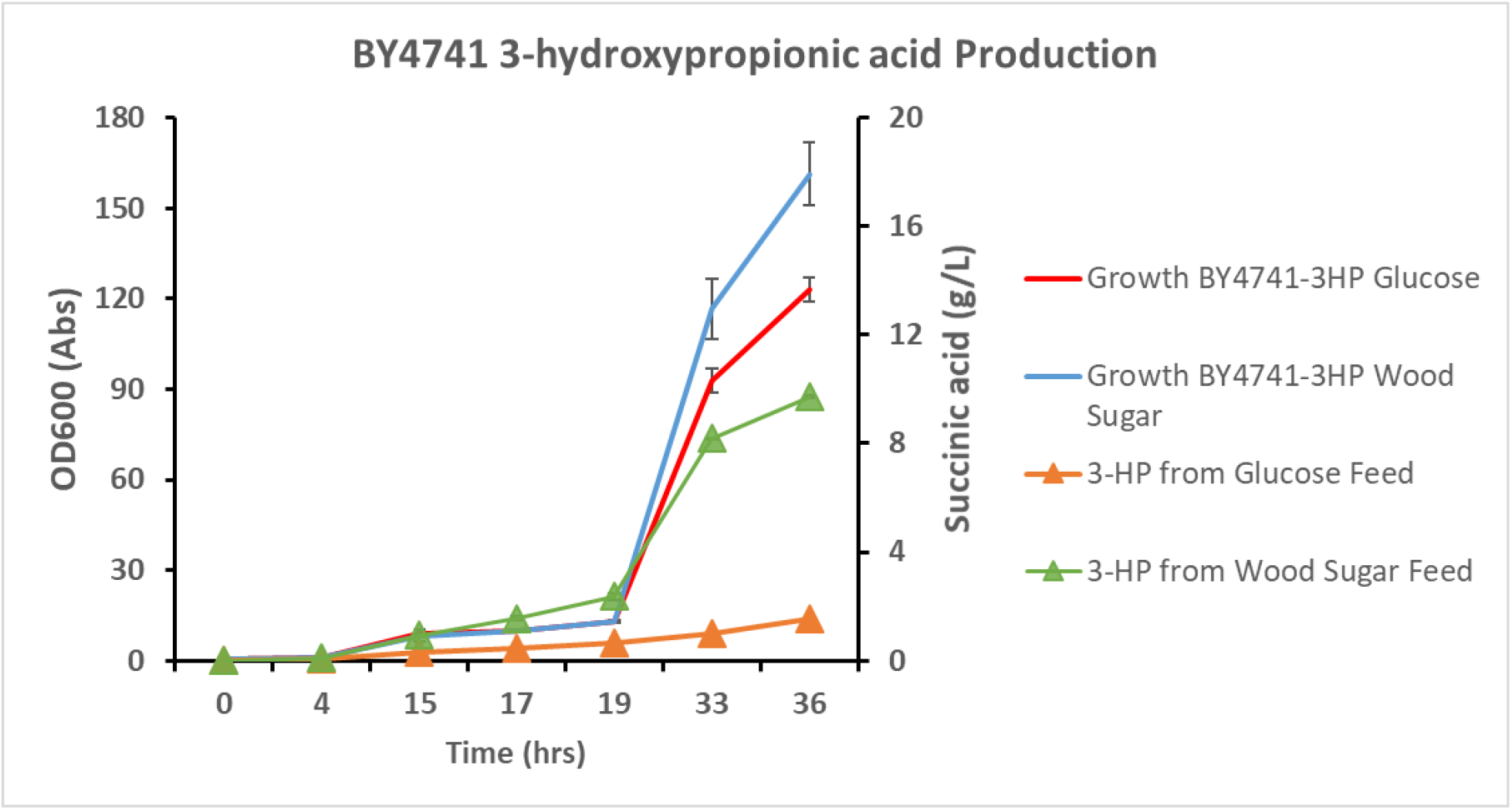
3-hydroxypropionic acid-producing yeast strain. Fed-batch over 36 hours with 300 g/L of either Glucose or Glucose-rich wood sugars. Error bars represent standard deviation. Each cultivation was carried out in either duplicates or triplicates.

**Figure 2.**
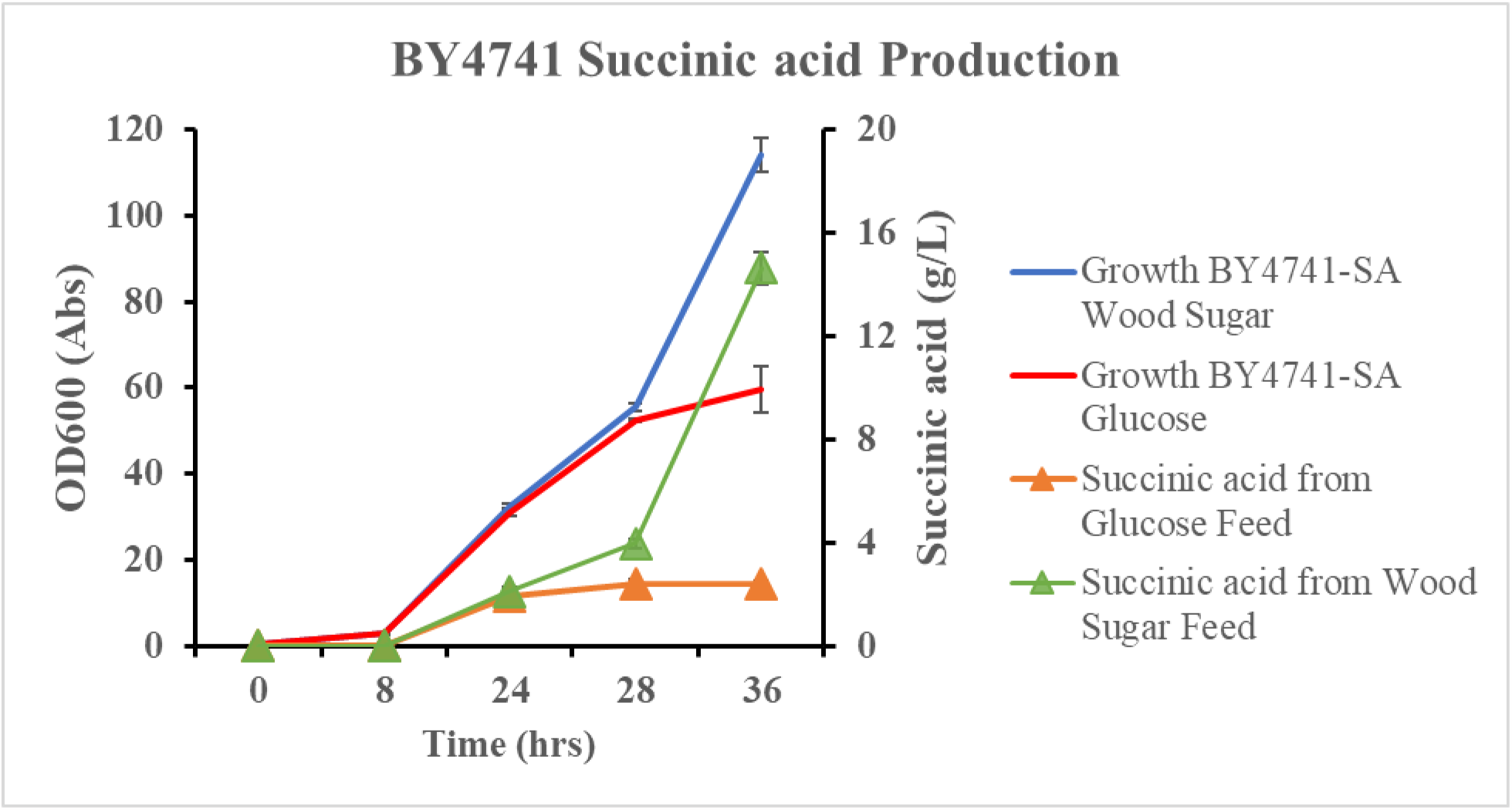
Succinic acid-producing yeast strain. Fed-batch over 36 hours with 300 g/L of either Glucose or Glucose-rich wood sugars. Error bars represent standard deviation. Each cultivation was carried out in either duplicates or triplicates.

## Discussion

In this study, utilizing exponential fed-batch cultivation strategies with glucose-rich wood sugars proved crucial for achieving substantially higher yields of 3-HP or SA than pure glucose feed. The maximum specific growth rates during the batch phase were 0.28 1/h (3-HP strain) and 0.295 1/h (SA strain). Towards the end of the batch phase, a detectable amount of ethanol, reaching up to 9 g/L, was observed, accompanied by increased respiration. This observation suggested the occurrence of the Crabtree effect, where carbon was being directed towards ethanol production, resulting in low biomass.

Upon lowering the maximum specific growth rate (μmax) to 0.15 1/h during carbon-limited feeding (Fig. 3), concentrations of organic acids (3-HP, SA, and some acetic acid, not shown) increased over time. This led to the initiation of uncoupling respiration from fermentation, aligning with findings from studies by Postma et al. (1989) and Verduyn et al. (1984). Maintaining the growth rate as low as possible under these conditions was crucial, facilitating CO_2_ production rates to match O_2_ uptake, thereby maximizing biomass yield.

**Figure 3.**
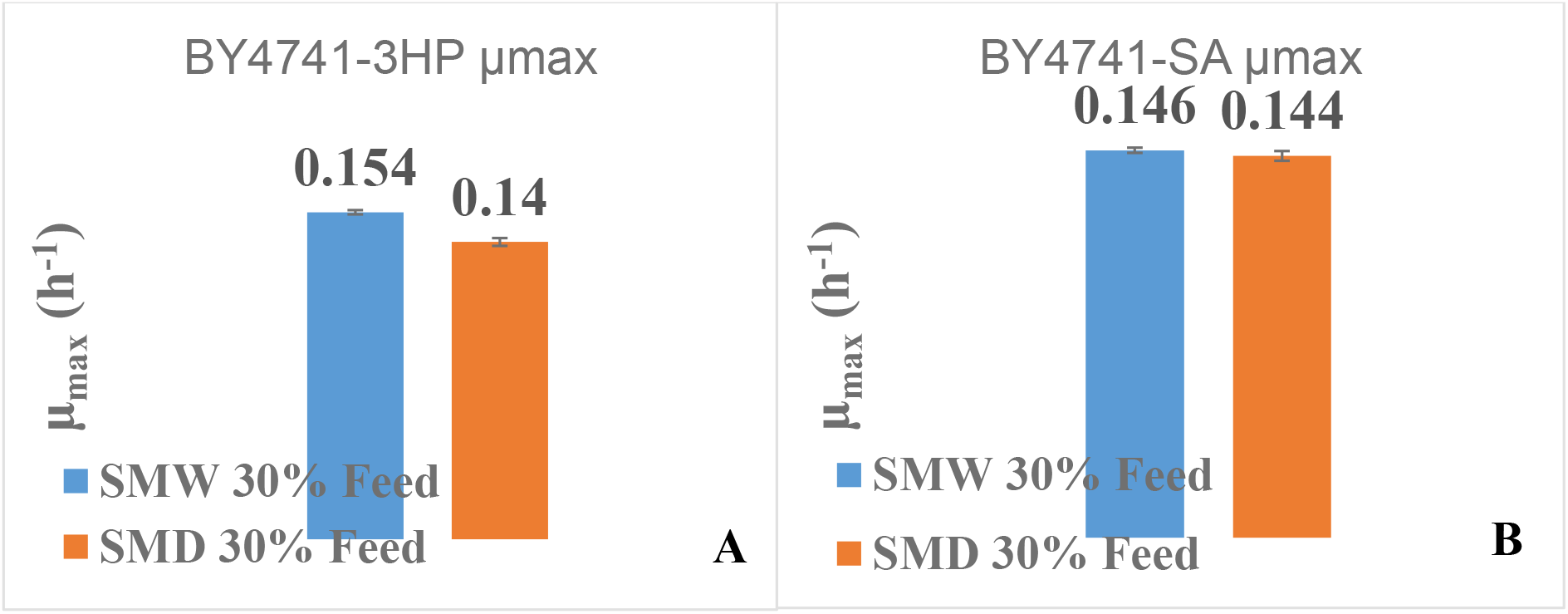
During the carbon-limited fed-batch phase, the target max specific growth rate was 0.15 1/h. A) For the 3-HP production strain, μ of 0.154 and 0.14 1/h were achieved for wood sugars and glucose growth, respectively. B) Similar results were attained with the succinic acid production strain.

Since 3-HP is a non-metabolizable organic acid for *S. cerevisiae* and SA production accumulates rapidly, the increasing levels of organic acids in the culture broth lowers the growth rate threshold at which ethanol formation occurs (Hagman et al., 2014). This underscores the importance of carefully managing growth conditions to optimize the production of target compounds in the fermentation process.

Numerous challenges arise in recovering 3-hydroxypropionic acid (3-HP) in high concentrations and purity. When microbial cell factories produce 3-HP, it completely solubilizes in culture medium at relatively low concentrations. Complicating matters, 3-HP is a carboxylic acid with a low molecular weight, enhancing its solubility in aqueous media. Traditional distillation techniques prove ineffective due to 3-HP’s decomposition at high temperatures and its tendency to polymerize at higher concentrations. The subsequent processing steps necessary for the recovery of purified 3-HP can represent a considerable portion of the total production costs (Chemarin et al., 2019). Specifically, this percentage can vary for carboxylic acids but remains substantial, explaining why 3-HP is commercially available as a 30% aqueous solution at a premium price from several chemical producers.

## Conclusions

This work shows that the wood-derived glucose-rich hydrolysate feeding composition produces the highest 3-HP and SA titers compared to glucose feeding of the same concentrations with engineered *S. cerevisiae* in this study. It is unclear why the wood sugars provide a better environment for 3-HP and SA production in yeasts, but similar results have been obtained in a recent study (Ingham *et al*., 2023), making the combination of high concentration of growth salts, wood sugars, yeast cell factories, low pH, and a slow feeding rate for 3-HP or SA production worth a more in-depth study.

## Declarations

Ethics approval and consent to participate

Consent for publication

Availability of data and material

Competing interests

Funding

ERDF and the Estonian Research Council supported this study via projects RESTA6

Authors’ contributions

Authors’ information

